# A system-level gene regulatory network model for *Plasmodium falciparum*

**DOI:** 10.1101/2020.08.10.235317

**Authors:** ML Neal, L Wei, E Peterson, ML Arrieta-Ortiz, SA Danziger, NS Baliga, A Kaushansky, JD Aitchison

## Abstract

Many of the gene regulatory processes of *Plasmodium falciparum*, the deadliest malaria parasite, remain poorly understood. To develop a comprehensive guide for exploring this organism’s gene regulatory network, we generated a system-level model of *Plasmodium falciparum* gene regulation using a well-validated, machine-learning approach for predicting interactions between transcription regulators and their targets. The resulting network accurately predicts expression levels of transcriptionally coherent gene regulatory programs in independent transcriptomic data sets from parasites collected by different research groups in diverse laboratory and field settings. Thus, our results indicate that our gene regulatory model has predictive power and utility as a hypothesis-generating tool for illuminating clinically relevant gene regulatory mechanisms within *Plasmodium falciparum*. Using the set of regulatory programs we identified, we also investigated correlates of artemisinin resistance based on gene expression coherence. We report that resistance is associated with incoherent expression across many regulatory programs, including those controlling genes associated with erythrocyte-host engagement. These results suggest that parasite populations with reduced artemisinin sensitivity are more transcriptionally heterogenous. This pattern is consistent with a model where the parasite utilizes bet-hedging strategies to diversify the population, rendering a subpopulation more able to navigate drug treatment.

## Introduction

Despite decades-long eradication campaigns, malaria remains a global burden with an estimated 405,000 deaths worldwide in 2018 (1), largely as a result of the deadliest malaria parasite, *Plasmodium falciparum* (*P. falciparum*). While malaria-associated deaths have declined over the past decade, likely due to both vector control measures and the roll-out of artemisinin-based combination therapies, deaths per year have plateaued in recent years. The origin of all malaria-associated mortality and morbidity is the destructive, cyclic asexual development of blood stage parasites that leads to erythrocyte death. Thus, elucidating the parasite’s molecular regulatory mechanics during its blood stage provides opportunities for identifying drug targets that would reduce the global burden of the disease. While extensive gene regulation at the level of translational repression occurs during the parasite’s mosquito-to-man and man-to-mosquito transitions (2–4), it appears that transcriptional, not translational regulation, plays a dominant role in protein regulation during the asexual, blood stage cycle (5). Transcriptional profiling has illustrated tight oscillatory patterns in cohorts of genes (6), suggesting that tightly regulated transcriptional networks initiate and/or respond to parasite life cycle progression within the asexual blood stage. However, outstanding questions that surround transcriptional regulation in *P. falciparum* asexual stages remain. To elucidate the gene regulatory interactions that contribute to key cellular processes in blood stage *P. falciparum*, we aimed to build a predictive genome-scale transcriptional regulatory network (TRN) for the parasite that could be applied to laboratory and field-isolated *P. falciparum* strains alike.

TRNs link transcription factors (TFs) with their targets and are generally constructed using a set of transcriptomes and a pre-existing set of TF-target pairs (7). The Apicomplexan Apetala2 (ApiAP2) family of DNA binding proteins is the dominant set of characterized TFs for *Plasmodium* (10), and has been demonstrated to bind specific DNA sequences (8) and regulate multiple steps in parasite life cycle progression (9, 10) including erythrocyte invasion (11), blood stage replication (10), sexual differentiation (12), oocyst development (13), sporogony (14) and liver stage development (15). Other TFs have been molecularly analyzed (16) and putative TFs have been compiled using computational approaches (17). Building a *P. falciparum* TRN allowed us to predict and quantify, on a genome scale, which of these transcriptional regulators influence which target genes, and to what extent.

To construct the model, we employed a set of system biology tools and an analysis methodology previously developed for constructing <u>E</u>nvironmental and <u>G</u>ene <u>R</u>egulatory <u>I</u>nfluence <u>N</u>etworks (EGRINs) in microbes such as *Halobacterium salinarum* (18, 19), *Escherichia coli* (18), *Mycobacterium tuberculosis* (20), and *Saccharomyces cerevisiae* (21) (Figure 1). We built the *P. falciparum* EGRIN (*Pf*EGRIN) by analyzing transcriptomic data sets to discover co-regulated groups of genes, and then performed regularized regression to determine which proteins regulate those genes, and to what extent. The result is a set of weighted interactions between transcription regulators (TRs) and their targets comprising a genome-wide regulatory network that can be used to predict target gene expression levels based on expression levels of their regulators (Figure 1).

**Figure 1.**
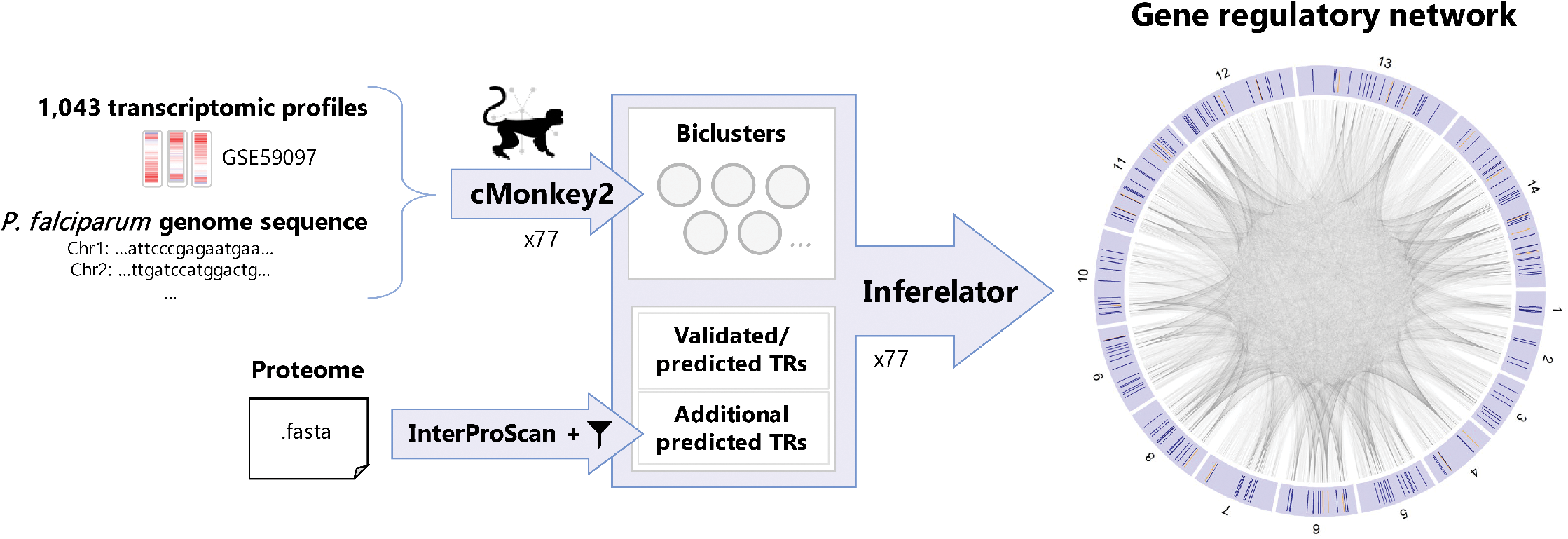
Overall workflow used to generate the global *P. falciparum* EGRIN. We used cMonkey2 to generate biclusters containing genes showing coherence across sample subsets, then used the Inferelator tool to combine that information with a list of transcription regulators (TRs) to generate a genome-scale regulatory network. A higher-resolution image of the network visualization on the right is provided in Figure S4. Network visualization generated using the “circos” R package (58).

In addition to outputting weighted TR-target interactions, EGRIN models facilitate the identification of concordant sets of transcripts associated with a given phenotype. One of the critical phenotypes in the context of malaria is the loss of artemisinin sensitivity. Artemisinin and its derivatives are widely used in combination therapies against malaria to quickly reduce the patient’s parasite biomass. Several countries have seen the emergence of parasite strains that require extended clearance times following artemisinin-based therapy, threatening one of the cornerstones of malaria treatment across the globe (22). Importantly, although genetic correlates of artemisinin resistance have been identified (23), including mutations in the Kelch13 protein (24, 25), not all parasites with a reduction in sensitivity of artemisinin harbor Kelch13 mutations, and the molecular pathways underpinning artemisinin resistance include global processes such as production of phosphatidylinositol 3-phosphate containing vesicles (26), oxidative stress (27), the unfolded protein response (28) and hemoglobin endocytosis (29). A comprehensive view of how a parasite is able to circumvent artemisinin has yet to be fully defined (30). To elucidate properties associated with artemisinin resistance, we identified concordant gene sets generated for the EGRIN that were associated with transcriptomic samples collected prior to artemisinin-based therapy that showed an artemisinin-sensitive (AS) or artemisinin-resistant (AR) phenotype after therapy. We hypothesized that molecular mechanisms underlying artemisinin resistance would manifest as gene sets showing elevated coherence among samples with longer parasite clearance times. Surprisingly, our analysis revealed that artemisinin resistance is linked to dramatically *less* coherence across a range of gene regulatory mechanisms, including – but not limited to - those associated with processes known to be part of the parasite’s bet-hedging strategy for evading the immune system during blood stage infection and persisting in the presence of environmental stress (31, 32).

## Materials and Methods

### Data sources

Transcriptomic data used to train our *Pf*EGRIN model were downloaded from Gene Expression Omnibus (GEO) accession GSE59097 using the R package GEOquery (33). This data set is associated with a study by Mok, et al. (28) that examined transcriptomic correlates of artemisinin resistance. It consists of microarray-based gene expression measurements on 1,043 *P. falciparum* field isolates obtained in Bangladesh, Democratic Republic of Congo, Cambodia, Laos, Myanmar, Thailand, and Vietnam. Each transcriptomic profile in this data set represents an individual patient’s infection, was obtained prior to artemisinin-based therapy, and is associated with an infection-specific parasite clearance half-life value. AR and AS parasites were found in all geographic regions where sampling occurred.

The Mok, et al. study also includes a separate 110-sample transcriptomic data set (GEO accession GSE59098) consisting of microarray measurements on blood stage parasites that were collected from 19 of the Pailin, Western Cambodia patients in accession GSE59097, cultured *ex vivo*, and then sampled at various timepoints over 40 hours. This data set was one of three used to assess the model’s ability to predict gene expression levels in transcriptomic data not used to train the model. The other two data sets used in this assessment are available through GEO accessions GSE83667 and GSE116341. GSE83667 consists of 58 microarray-based, *P. falciparum* transcriptomic profiles obtained from blood samples collected in Malawi (34). GSE116341 primarily consists of single-cell RNA-seq data from an *in vitro P. falciparum* blood stage strain, but also includes 28 bulk RNA-seq measurements on parasite populations, which we used for model validation (35).

FASTA and GFF files containing genome sequence and annotation information for the *P. falciparum* 3D7 strain were downloaded from plasmodb.org (36) (release 42) and used in cMonkey2 runs.

A FASTA file containing *P. falciparum* 3D7 protein sequence information across the organism’s proteome was downloaded from UniProt (37) for use in identifying proteins with potential gene regulatory functions.

### Model construction

To create our global gene regulatory network, we adapted a workflow previously used to generate EGRINs for *Halobacterium salinarum* and *Escherichia coli* (18). First, we used the cMonkey2 biclustering tool (38) (version 1.2.11) to identify co-regulated genes among the transcriptomic profiles from training data (GEO accession GSE59097). We then used the Inferelator (19, 39) (https://github.com/baliga-lab/cMonkeyNwInf) to generate TR-target networks from the collection of biclusters in each cMonkey2 run in combination with a list of TRs. TR-target weights were averaged across these EGRINs to create an ensemble EGRIN. The final list of 90,300 TR-target pairs comprising the model was determined by ranking all pairs in the ensemble EGRIN by their absolute weights, then testing model predictions on training data using various percentages of the top-ranked pairs.

### Identifying biclusters from transcriptomic data

To prepare the transcriptomic data from the GSE59097 training data set for cMonkey2 biclustering analysis, we normalized the data on a gene-by-gene basis using a z-score transform. In accordance with established cMonkey2 protocols, and for comparability to previous EGRIN development efforts, the 5,000 genes with the highest transcript variance were used in our runs. Runs were performed using the Bicluster Sampled Coherence Metric (BSCM), which helps optimize the difference in coherence between samples within a bicluster and those not in the bicluster (40). cMonkey2 runs were performed with *de novo* binding motif detection enabled. This means that when the tool calculates the likelihood that a gene belongs within a particular bicluster, it scans the *P. falciparum* genome to determine if the gene and those in the bicluster share common upstream binding motifs and are therefore more likely to be co-regulated.

### Determining bicluster convergence

To determine the number of cMonkey2 runs needed to ensure robustness in our final gene regulatory model, we performed multiple cMonkey2 runs, then used separate Inferelator runs to compute TR-target weights for each individual cMonkey2 run. We determined what percentage of TR-target pairs produced by the Inferelator had average weights that shifted less than 5% when comparing average weights over the total number of cMonkey2 runs to average weights over three less than the total number of runs. If more than 95% of the averaged TR-target weights shifted less than 5%, then we concluded that additional cMonkey2 runs would not substantially change the gene regulatory network model derived from the averaged weights. We reached our target level of convergence for TR-target interaction weights after 77 cMonkey2/Inferelator runs: less than 5% of the average weights on TR-target interactions changed more than 5% between 74 and 77 runs (Figures S1 and S2).

### Compiling *P. falciparum* transcription regulators

The list of TRs used as input to the Inferelator consists of known, empirically-analyzed TFs, predicted TRs reported in the literature, and predicted TRs we identified through a protein domain analysis performed on the *P. falciparum* proteome. In this context, when discussing the transcriptional influence of gene products on other genes’ expression, we make a distinction between TFs and TRs. We refer to proteins that regulate target gene expression by binding to *cis*-regulatory DNA elements as TFs. We consider TRs a broader set of proteins that regulate gene expression, but not necessarily through sequence-specific DNA-binding. We make this distinction because many of the proteins presumed to have gene regulatory activity in our model have not been molecularly analyzed, and their precise mode of action as a regulator remains unknown.

The TR list was populated in part using the collection of regulatory genes previously compiled by Bischoff and Vaquero (17) who used Pfam Hidden Markov Model profiles (41) to identify proteins with potential transcriptional activity. Among the 202 genes they identified, their list includes the 27 genes encoding ApiAP2 proteins (42). We also included empirically-analyzed and putative TFs mentioned in a recent review (16). To further expand the list, sequences for proteins in the *P. falciparum* proteome were downloaded from UniProt (37) and then input to the InterProScan tool (43) to identify protein domains within each proteome entry. InterProScan utilizes domain information from several databases, including Pfam (41) and SMART (44), and provides textual descriptions of any domains that are found within the scanned proteins. From the InterProScan output, we selected any proteins annotated against protein domains containing the phrase “DNA binding” or “transcription factor”. We accounted for subtle variations in these specific phrases. For example, domain annotations containing the phrase “dna-binding”, “DNA binding” or “dna_binding” were all flagged by our approach. We added the genes encoding these proteins to the existing list of TRs, accounted for overlap, then selected those genes that were present in the list of 5,000 used for our cMonkey2 runs. This final list of genes was used as input to each of the Inferelator runs we performed to generate our ensemble of EGRIN models.

### Model evaluation

Evaluations using the three transcriptomic validation data sets compared measured and model-predicted mean gene expression across biclusters generated from our cMonkey2 runs. For all three validation data sets, the model was used to predict expression values for each gene in a bicluster using the measured expression levels of the TRs regulating those genes and the weight of those interactions in our model. For each bicluster tested, the mean model-predicted expression of its genes was computed and compared to the mean measured expression from the validation data set. To ensure the comparisons were valid, we normalized the validation data using the same method applied to the training data used to create the model (gene-by-gene z-score transform). To quantify model accuracy, we calculated root-mean-square errors (RMSEs) between the predicted and measured values across samples in each bicluster. In some cases, due to missing values in samples within the validation sets, we were not able to compare the full complement of predicted and measured values for each sample. Therefore, for a bicluster to be included in this analysis, we required that more than half of the samples in the validation data set contain the full complement of predicted-to-measured gene expression comparisons. An archive of R scripts and data objects that allow a user to reproduce model-based, quantitative predictions reported here is included as Supplementary Data.

For evaluations that assessed overlap between model-predicted targets of ApiAP2 TFs and targets predicted based on empirically-derived binding sequence motifs, we used the “TF Binding Site Evidence” page on plasmodb.org to retrieve lists of genes that are targeted by ApiAP2 TFs based on the presence of TF-specific upstream binding motifs. We used the page’s default parameters to set the size of the upstream region to scan (1000 bp), the minimum number of motifs per gene (1 motif) and the minimum confidence level for a match to the motif (*P*-value ≤ 1e-4). We then compared the retrieved lists to the targets of ApiAP2 TFs predicted by our model. For each ApiAP2 TF evaluated, a hypergeometric enrichment test was used to quantify the overlap between motif-based and model-based target predictions.

### Identifying correlates of artemisinin resistance

Hypergeometric enrichment tests were used to identify biclusters that were either significantly enriched for samples with AR infections or significantly enriched for samples with AS infections. In accordance with previous criteria used to analyze the training data used to build our model, we classified samples with parasite clearance half-lives greater than or equal to 5 hours as AR (28). Samples with lower clearance times were classified as AS. After identifying biclusters that showed over-representation of either the AR or AS group, we functionally profiled the genes in the biclusters by determining their enrichment for members of gene sets collected from the Gene Ontology (45, 46), the Malaria Parasite Metabolic Pathways (MPMP) resource (47, 48), and data sets associated with stage-specific gene expression at different points in the parasite life cycle (49–56).

## Results

### Gene Regulatory Network Model Construction

To generate a robust *P. falciparum* EGRIN, we adapted a workflow previously used to create EGRINs for *Halobacterium salinarum* and *Escherichia coli* (18). As illustrated in Figure 1, we first used the biclustering tool cMonkey2 (57) to identify genes and transcriptomic samples that group together based on gene expression coherence. These biclusters were computed based on a large *P. falciparum* transcriptomic data set (GEO accession GSE59097) and the genomic sequence of the organism. We then used the Inferelator tool (19, 39) to generate a network of TR-target interactions based on coherence between TRs and biclustered genes. A list of TRs was compiled from known, empirically-analyzed TFs (16), putative regulators compiled via computational methods in a previous study (17), and an analysis we performed using InterProScan (43) that identified additional proteins with potential regulatory activity. We reserve the TF term for proteins that bind to *cis*-regulatory DNA elements; TRs are a superclass of proteins whose regulatory activity is not necessarily limited to sequence-specific binding. In the following sections, we detail results from each step in our workflow which ultimately culminated in our system-level gene regulatory network.

### Generating biclusters

The cMonkey2 biclustering tool identifies potential gene regulatory programs based on transcriptomic data and *a priori* biological knowledge (38). The tool groups genes based on their *coherence* across transcriptomic profiles; genes with expression levels that shift more in parallel across transcriptomic profiles are more likely to be grouped together by the algorithm as this suggests the genes may be under the influence of a common set of transcriptional regulators. Concomitantly, cMonkey2 identifies subsets of transcriptomic profiles (samples) in which gene sets show high coherence. This allows cMonkey2 to delineate particular experimental samples in which gene coherence is high. For example, expression in a certain regulatory pathway may only be coherent among transcriptomic profiles from samples that were treated with a certain drug. The primary product of a cMonkey2 run, therefore, is a set of biclusters, each of which contain a set of genes and a set of samples whose transcriptomic profiles show elevated coherence among those genes. These biclusters identify genes that may be part of a common regulatory program and thus, form the foundation of EGRIN models (Figure 2). The difference between coherent gene expression among samples within a bicluster and the remaining samples in the data set can be visualized by normalized gene expression (Figure 2, top) or by examining the standard deviation between transcript levels of genes within the bicluster (Figure 2, bottom).

**Figure 2.**
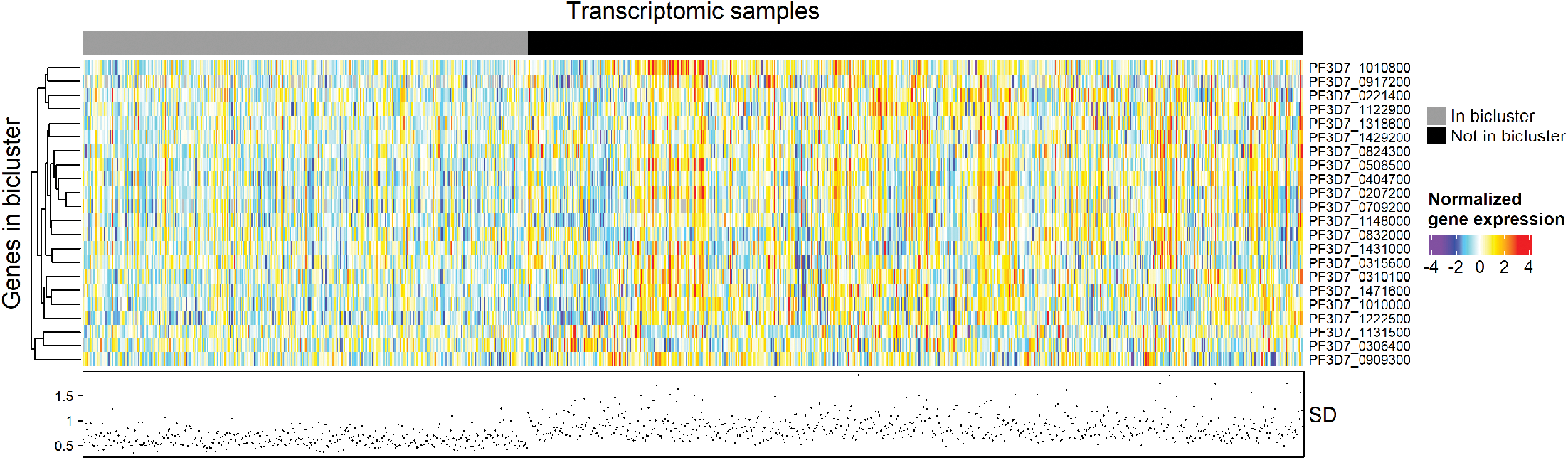
Example of cMonkey2 biclustering. Heatmap colors indicate normalized expression values for genes in an example bicluster across 380 transcriptomic samples within the bicluster (gray annotation bar) and 663 samples not in the bicluster (black anno tati on bar). Scatterplot below heatmap indicates standard deviation (SD) of expression across genes for each sample. Samples in the bicluster show lower SD (more coherence) compared to the samples not in the bicluster.

The cMonkey2 algorithm includes elements of randomness as it attempts to optimally group genes and samples together. For example, at the beginning of each cMonkey2 run, genes and samples are randomly divided into a specified number of biclusters. Additionally, to avoid falling into local minima, a small amount of random noise is applied to the probability matrix used to assign genes to biclusters at each iteration of the algorithm. At the end of each iteration, a random set of elements (genes or samples) is selected for bicluster re-assignment as part of the algorithm’s optimization routine as well (18). Therefore, separate cMonkey2 runs using the same input data do not generate identical results, and an ensemble of cMonkey2 runs is needed to account for run-to-run variation and produce statistically robust results. To determine the required number of cMonkey2 runs, we performed runs until 95% of the TR-target weights generated by the Inferelator from each run showed convergence based on their average values across runs. We found that our model showed convergence after 77 runs (Figures S1, S2).

### Identifying candidate transcription regulators

Building an EGRIN model requires both cMonkey2 runs and a list of TRs as input to the Inferelator (Figure 1). Our compiled list of TRs includes the ApiAP2 proteins, a previously compiled list of computationally-predicted transcription-associated proteins (17), empirically-analyzed and additional candidate TRs (16), and additional proteins that we identified as having potential regulatory activity. We identified this latter group of proteins in the interest of being inclusive in our TR list, given that much of the gene regulatory landscape of *P. falciparum* remains uncharacterized. Using the InterProScan tool, we identified proteins in the *P. falciparum* proteome containing protein domains that indicate potential gene regulatory functions. Out of the full *P. falciparum* proteome, 4,996 proteins were annotated with at least one protein domain. Of these, 105 were annotated with domains that suggested a potential role as a TR, 92 of which were in the list of 5,000 genes used as input to the cMonkey2 runs (see *Identifying biclusters from transcriptomic data*). Twenty-four of the 92 were in the list of TRs that we had already compiled from literature sources. Thus, through this analysis, 68 novel candidate TRs were added to the previously compiled list of TRs, bringing the number of TRs used to train our EGRIN to 258.

### Generating and quantifying the gene regulatory influence network

Using a set of TRs and the biclusters from a cMonkey2 run, the Inferelator uses elastic net regularization, which is a linear regression-based statistical modeling method, to produce a list of TR-bicluster pairs with weights that quantify the influence of the TR on the genes in the bicluster (Figure 1). These weights are then assigned to each gene in that bicluster to generate a network consisting of weighted TR-target gene pairs. For each of our cMonkey2 runs, we performed a separate Inferelator run to generate a TR-target network, then aggregated the results from the Inferelator runs into an ensemble network using the approach established by Brooks, et al. (2). Using an input set of TR expression values in the ensemble network, we can then predict the expression level of an individual gene by identifying which TRs regulate the gene, then computing the dot product of the weights on the TRs regulating the gene and the TRs’ expression levels.

The Inferelator identifies regulatory relationships between genes based on correlations in expression levels; thus, one of the challenges in developing a network from a set of correlations is determining the directionality of regulatory relationships. For example, if two TRs have well-correlated expression levels, this may indicate that the first regulates the second, the second regulates the first, or the regulation is reciprocal. If given temporally-resolved expression data as input, the Inferelator can address this issue using dynamic modeling of TR-target interactions. The nature of the sampling here precluded the incorporation of temporally resolved data. The data used for model training was from one time point, and dynamic modeling was therefore not applicable. However, an additional way to address the directionality issue, which was used in our approach, is by applying *de novo* upstream motif detection when using cMonkey2 to identify biclusters. Doing so adds an additional layer of biological evidence indicating that co-regulated genes in a bicluster are targets of some TR and not necessarily the reverse. Thus, while the Inferelator may find a regulatory relationship suggesting that TR1 targets TR2, it may not necessarily predict the reciprocal because upstream motif detection may assign TR1 to a bicluster whose overall expression profile does not correlate well with TR2. We therefore assume that each individual TR-target regulatory relationship identified by the Inferelator is unidirectional. Reciprocal relationships may nonetheless appear in the resulting network if the Inferelator separately identifies both elements of a reciprocal relationship between TRs.

In accordance with previous EGRIN modeling efforts, we initially used the top 100,000 TR-target pairs with the highest absolute weights in our ensemble network for our *Pf*EGRIN model. To determine if this cutoff would produce the most predictive model, we performed an optimization analysis that compared the predictive capabilities of models consisting of different numbers of top-ranked TR-target pairs. This analysis assessed the accuracy of model predictions on mean bicluster gene expression across the 38,500 biclusters generated by our 77 cMonkey2 runs. We found that a model consisting of 7% (90,300) of the top-ranked TR-target interactions from the ensemble gene regulatory network provided the best fit to the training data based on mean RMSE values (Figure S3). The final *Pf*EGRIN model, therefore, consists of these 90,300 TR-target pairs. An R data object file listing these interactions and their weights is provided in the Supplementary Data.

### Gene regulatory network characteristics

The final *Pf*EGRIN model consists of 5,000 genes (nodes) and 90,300 TR-target interactions (directed edges) (Table 1, Figure S4). Positive regulation of target genes by TRs dominates (86%) the network. All 258 TRs are regulated by at least one other TR. Five TRs had no influence on the expression of other genes in the model but were regulated by other TRs. Thus, the model includes 253 TRs that influence the expression of target genes.

**Table 1.**
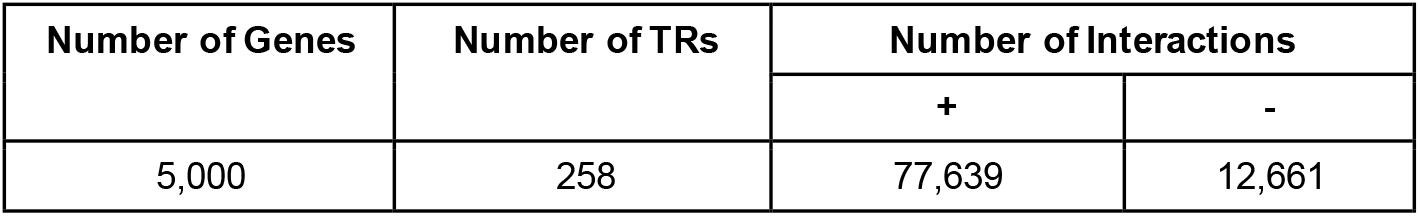
Connectivity metrics from the TR-target regulatory network.

To investigate the topology of the TR-gene regulatory network, we performed a network analysis focused on outgoing connectivity of TR nodes. The majority of TRs regulate a small number of target genes, while few TRs regulate a relatively large number of target genes (Figure 3A). Comparing the probability density of TRs and their out-degrees on a log-log scale revealed a linear relationship (Pearson correlation coefficient = −0.86, *P*-value=3.6e-9), indicating that the network degree distribution follows a power-law (Figure 3A, inset). This suggests a scale-free topology, which is a general characteristic of many networks in nature and society (58–60). For a scale-free network, the probability of a node having an out-degree of *k* follows the power-law function

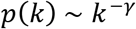

where γ is the power law scaling exponent. The scaling exponent of our *P. falciparum* gene regulatory network is 1.01, which is smaller than empirically-derived exponents obtained from other eukaryotic model organisms including *C. elegans* (4.12), *D. melanogaster*(3.04), *S. cerevisiae* (2.0) and *A. thaliana* (1.73) (61). The scale-free connectivity suggests that gene regulation is performed by a relatively low number of highly influential TRs. The comparatively small scaling exponent suggests that a larger fraction of genes is targeted by these highly-influential TRs (“hubs”) than in other organisms. In other words, the same number of highly-influential TRs in the *Pf*EGRIN target a disproportionately larger number of genes than in regulatory networks with higher scaling exponents.

**Figure 3.**
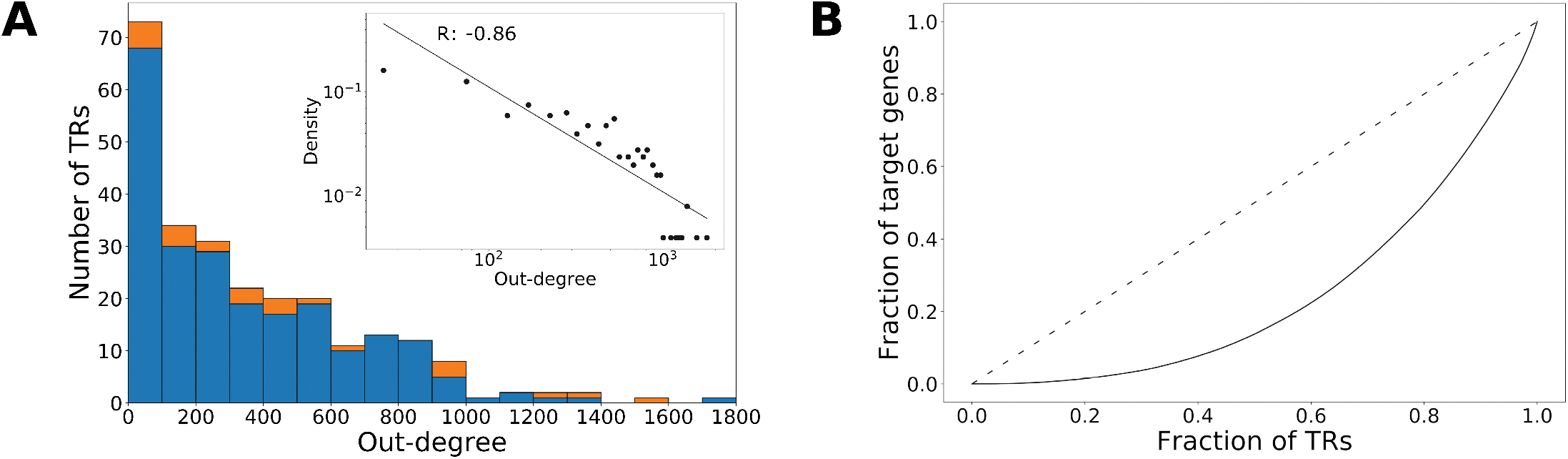
Outgoing connectivity of gene regulatory network. (**A**) Histogram of out-degree distribution. Orange shading in stacked bar plot indicates number of ApiAP2 TFs in a bin; blue shading indicates number of other TRs. Inset: Probability density of TRs and their out-degrees on a log-log scale with regression line. R is the Pearson correlation coefficient. (**B**) Lorenz curve of TRs and their target genes. Dashed line indicates line of equality.

The majority of TRs (249/258) included in the EGRIN model influence less than 1,000 genes. There are nine TRs that each influence more than 1,000 genes with PF3D7_0407600 having the highest out-degree (1,791). These nine TRs regulate 87.8% of the target genes (4,390). We ranked the TRs based on their out-degrees and plotted the cumulative proportion of TRs against the cumulative proportion of the corresponding target genes (Lorenz curve (62); Figure 3B). The deviation of the curve from the line of equality again demonstrates that a small fraction of TRs regulates a significant number of target genes in the EGRIN model.

### Model predictions on gene expression

We evaluated the capability of the *Pf*EGRIN to predict transcriptomic gene expression levels using three validation data sets not used to train the model. These transcriptomic data sets were selected so that the model would be tested against data obtained from a variety of research groups, using a variety of measurement modalities, and from blood-stage parasite populations originating from different regions (Table 2). This allowed us to assess the model’s predictive capabilities across geographic locations, across transcriptomic platforms/processing pipelines, and across parasite populations cultured from clinical isolates or from *in vitro* laboratory stocks.

**Table 2.** Descriptions of transcriptomic validation data sets used to evaluate the *Pf*EGRIN model’s predictive performance and summary statistics for each test. Summary statistics for root mean square error (RMSE) and correlation coefficients are presented as median values followed by the inter-quartile range.

|  | Validation test 1 | Validation test 2 | Validation test 3 |
| --- | --- | --- | --- |
| *GEO accession* | GSE59098 | GSE83667 | GSE116341 |
| *Associated publication* | Mok, et al. 2015 (28) | Milner, et al. 2012 (34) | Ngara, et al. 2018 (35) |
| *Number of samples* | 110 | 58 | 28 |
| *Source of parasites* | Clinical isolates (Cambodia) | Clinical isolates (Malawi) | *In vitro* strain Pf3D7 |
| *Measurement technique* | Microarray | Microarray | RNA-seq |
| *Biclusters evaluated* | 10,462 | 6,341 | 10,078 |
| *Unique genes evaluated* | 4,515 | 3,985 | 4,473 |
| *RMSE for PfEGRIN, training data* | 0.26 (0.21-0.44) | 0.28 (0.21-0.44) | 0.26 (0.21-0.44) |
| *RMSE for PfEGRIN, validation data* | 0.29 (0.22-0.44) | 0.29 (0.22-0.38) | 0.33 (0.25-0.44) |
| *RMSE for random model, validation data* | 0.73 (0.65-0.81) | 0.72 (0.64-0.81) | 0.53 (0.42-0.68) |
| *Correlation coefficient for PfEGRIN, training data* | 0.97 (0.96-0.98) | 0.98 (0.97-0.98) | 0.97 (0.96-0.98) |
| *Correlation coefficient for PfEGRIN, validation data* | 0.95 (0.92-0.97) | 0.94 (0.90-0.98) | 0.85 (0.69-0.92) |
| *Correlation coefficient for random model, validation data* | 0.16 (0.03-0.32) | 0.18 (0.03-0.34) | 0.13 (-0.03-0.29) |

Overall, we found that predictions on validation data were only slightly less accurate than predictions on the training data (Table 2). We evaluated the distribution of RMSE values (Figure 4A) and correlation R values (Figure 4B) when using the *Pf*EGRIN model to predict expression values for biclustered genes on samples from the training set (those within a bicluster) and on all samples from the validation set, or when using a model that predicts expression based on random value selection from the normal distribution. Figure 5 shows sample-by-sample model predictions on validation data set 1 (GEO accession GSE59098) for an example bicluster in which predictive accuracy on the validation set was equal to the median of the model’s overall performance and is representative of all biclusters tested. Figure 5A compares, across samples in validation data set 1, the measured mean normalized gene expression values for genes in the bicluster against model-predicted values. Figure 5B shows the correlation between these values. The RMSE and correlation coefficients for these predictions on validation data set 1 are similar to the performance of the model on the training data shown in Figures 5C and 5D.

**Figure 4.**
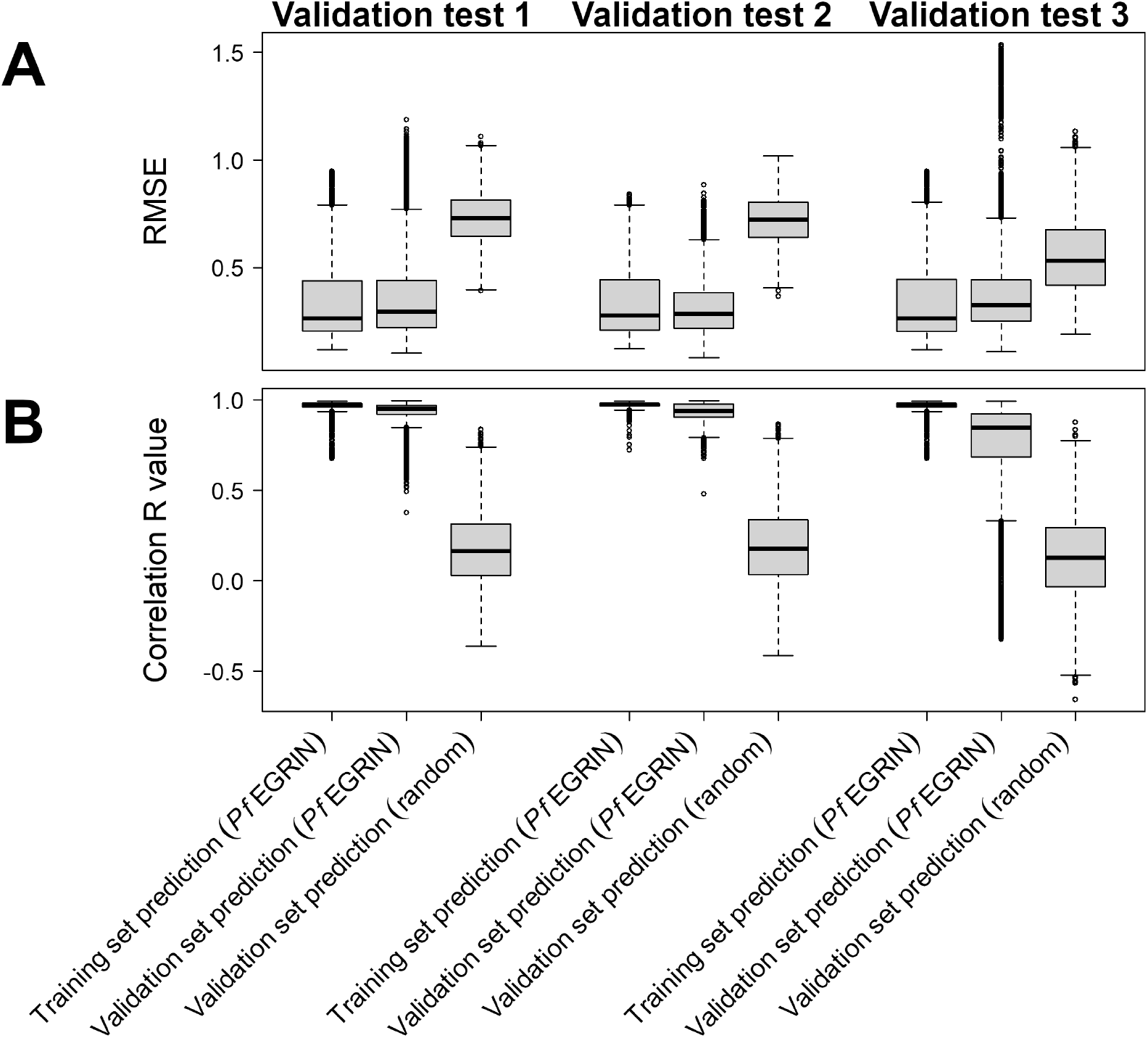
*Pf*EGRIN model predictive performance. (**A**) Distributions of root-mean-squared error (RMSE) in model-based gene expression predictions across cMonkey2 biclusters generated in building the model. For each of our three validation tests, we used the model to predict mean expression among biclustered genes across biclustered samples from training data as well as samples from validation data. For comparison, we also made predictions on the validation data using a model that randomly selects values from a normal distribution. (**B**) Pearson correlation coefficients (R) for the same predictionsin (**A**).

**Figure 5.**
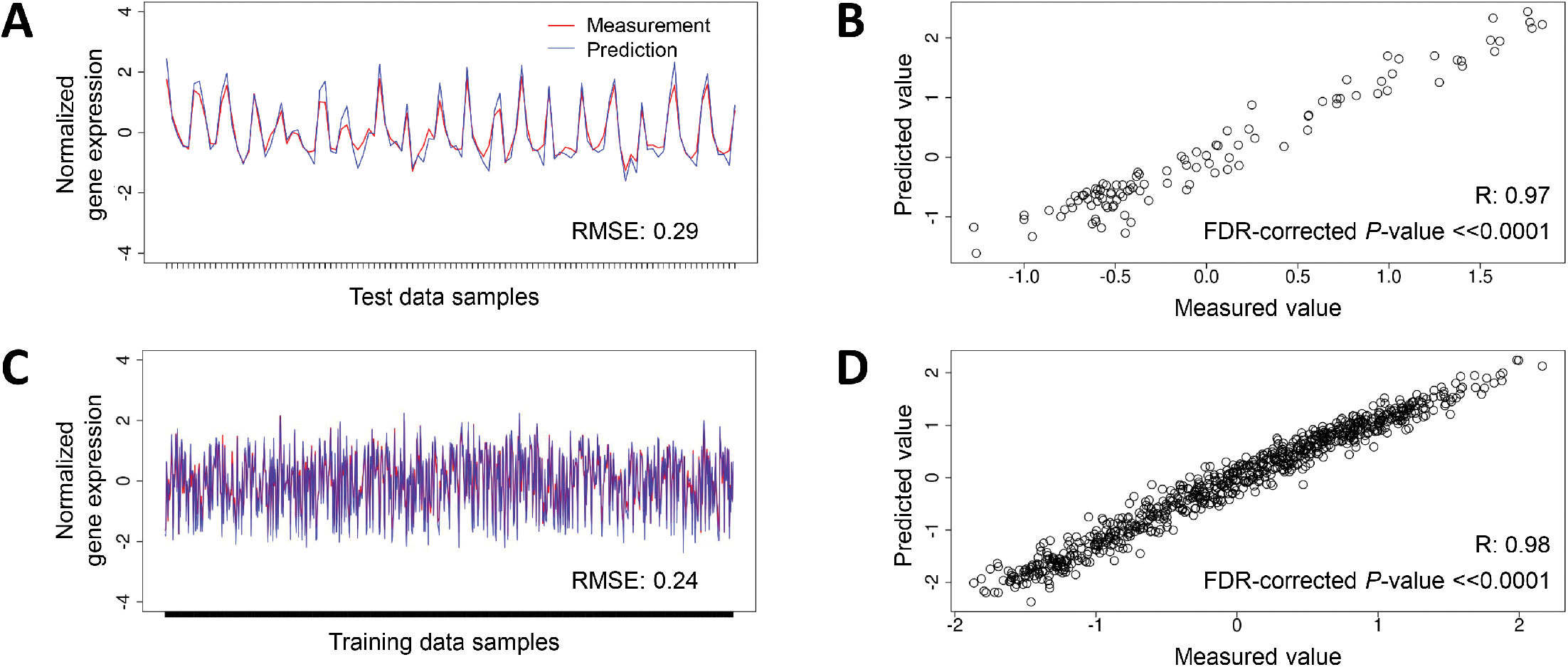
Representative predictions made by the *P. falciparum* EGRIN model. (**A**): Measured gene expression levels from the validation test 1 data set (red) compared to model predictions (blue) for genes in a representative cMonkey2 bicluster. Model accuracy in predicting expression levels for genes in this bicluster was similar to the model’s overall prediction performance for the validation set. (**B**) Scatterplot showing correlation between measured and predicted expression values in *A*. R is the Pearson correlation coefficient. (**C** and **D**): Same as in (**A**) and (**B**), with measured and predicted expression levels for samples in the GSE59097 training set contained in the bicluster.

The median RMSE values and correlation coefficients for model-based predictions on data from samples used to train the model were similar across our three validation tests (Table 2, Figure 4). Although similar, these values are not equivalent because the three validation sets had different complements of missing data. Thus, the fraction of the 38,500 biclusters generated by our 77 cMonkey2 runs that could be used for evaluation differed between the three validation tests, and the number of unique genes used in evaluating model predictions was also different between the tests.

Median RMSE scores were similar for *Pf*EGRIN predictions over the three validation tests and were significantly lower than those based on random predictions (*P* < 2.2e-16 for each validation set; paired Wilcoxon rank-sum test). Median correlation coefficients between *Pf*EGRIN-predicted and measured expression values were high (greater than or equal to 0.85) across the training and validation data sets (Table 2, Figure 4B).

### Model-based versus motif-based ApiAP2 target prediction

We next evaluated the capacity of the *Pf*EGRIN to identify targets of ApiAP2TFs, which regulate gene expression throughout different stages of the *P. falciparum* life cycle. We assessed the agreement between the model-predicted gene targets of ApiAP2 TFs and targets predicted based on empirically-determined upstream sequence motifs specifically recognized by those ApiAP2 TFs (8). Lists of predicted targets for each ApiAP2 based on upstream binding motifs were downloaded from plasmodb.org and compared to targets predicted by our model. Out of 18 ApiAP2 TFs compared, four showed significant overlap between motif-based and model-based target lists (hypergeometric test, FDR-corrected *P*-value < 0.05). In order of significance, these ApiAP2s were PF3D7_0802100, PF3D7_1456000, PF3D7_0604100, and PF3D7_1305200. Notably, mRNA abundances of the top three TFs have similar dynamics during the parasite’s intraerythrocytic developmental cycle (IDC), peaking in the early-mid schizont stage (8).

### Incoherent expression is associated with artemisinin resistance

Because the *P. falciparum* EGRIN was built using transcriptomes from parasites with various levels of artemisinin sensitivity, we reasoned that we might be able to identify biclusters that exhibit concordance selectively in samples showing AR. However, across all 38,500 biclusters generated in building the *Pf*EGRIN, no FDR-corrected *P*-values for AR sample enrichment fell below 0.81. In contrast, 3,202 biclusters (8%) met an FDR-corrected *P-*value cutoff of 0.05 for enrichment of AS samples (Figure 6). This suggested that AR is primarily associated with *in*coherence rather than coherence among gene regulatory programs.

**Figure 6.**
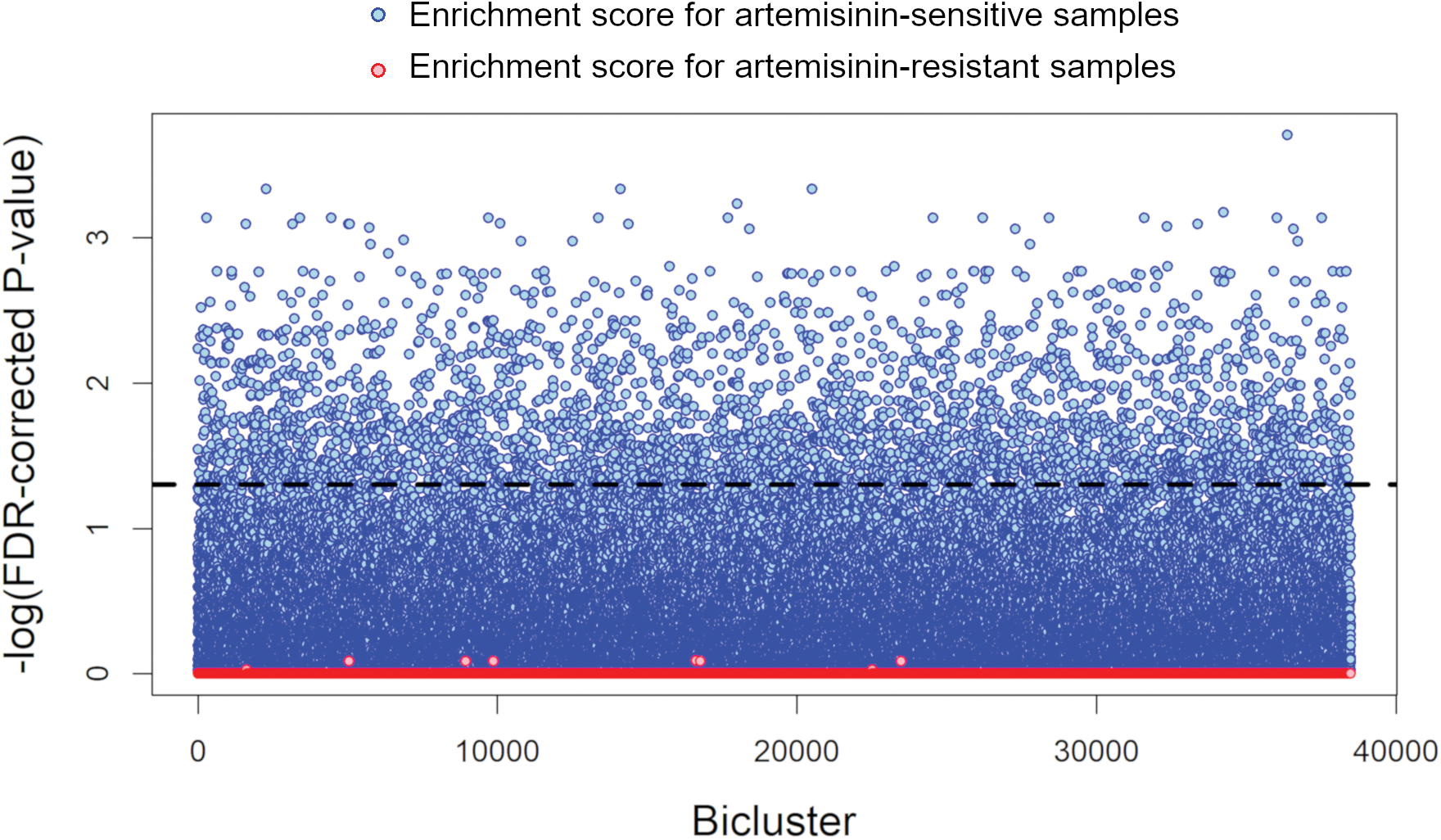
Hypergeometric enrichment scores for artemisinin-resistant and artemisinin-sensitive samples among all biclusters generated in constructing the *Pf*EGRIN. Dashed line indicates −log10 of an FDR-corrected *P*-value of 0.05.

### Correlates of artemisinin sensitivity

Considering gene expression was incoherent among AR samples, we identified the genes and biological functions most strongly associated with the AS biclusters. We first selected the most high-confidence AS biclusters by identifying those with FDR-corrected enrichment *P-*values less than 0.01. We then tabulated and ranked the number of times each gene appeared in one of these 710 highly enriched AS biclusters. We identified 111 genes that appeared more often than would be expected by chance (hypergeometric test, FDR-corrected *P*-values < 0.05) (Table S1). Among this set are 35 *var*, 27 *rifin*, 8 putative ribosomal component, 2 ApiAP2 TFs, 2 putative proteasome subunit, and 2 putative dynein heavy chain genes. Interestingly, we found that the highest-ranking genes were members of the parasite’s *var* gene family, which encode a group of immunovariant proteins involved in antigenic variation and erythrocyte adhesion at sites of vascular infection. They are a central component of the parasite’s strategy for evading immune clearance by the host, and a body of evidence suggests they are under the control of epigenetic pathways (reviewed in (63)); however the interpretation of this result based on microarray data is confounded by the high sequence complexity of genes families such as the *var* and *rifin* families (see Discussion).

To assess the broader landscape of biological functions associated with AS biclusters, we assessed their enrichment for various functional gene sets from GO, the MPMP resource, and sets associated with specific stages of the parasite life cycle. We identified 115 gene sets for which AS biclusters were significantly enriched (Figure 7, hypergeometric test, *P* < 0.05). For this analysis we initially defined enrichment scores with FDR-corrected *P*-values less than 0.05 as significant. However, a substantial fraction of AS biclusters (68%) show no enrichment across gene sets using this criterion and thus could not be functionally profiled. To help illuminate the functional roles of these biclusters, we relaxed the criterion and defined enrichment scores with uncorrected *P*-values less than 0.05 as significant. To ensure that we only included gene sets that showed enrichment more often than would be expected by chance, we determined the null distributions of the gene sets for our biclusters by randomly sampling 710 biclusters from our complete set of 38,500 and performing functional gene set enrichment tests on them. This process was repeated 10,000 times to build null distributions for each functional gene set. For our heatmap analysis, we only included those gene sets that appeared significantly more often in the 710 AS biclusters as compared to occurrences in their null distribution (95^th^ percentile or higher).

**Figure 7.**
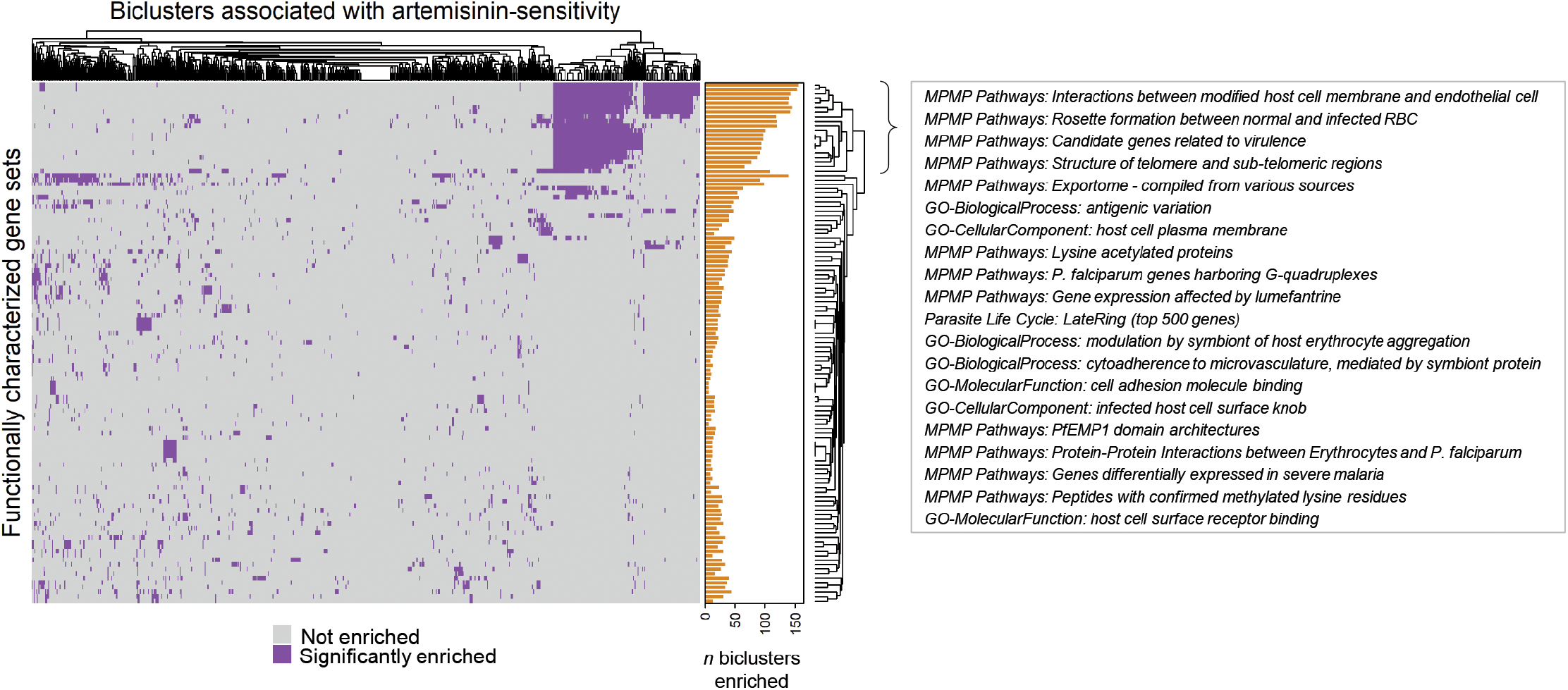
Heatmap showing clusters of functional gene sets among the 710 biclusters most highly enriched for artemisinin-sensitive samples. Annotation bar plot indicates the total number of biclusters that were enriched for a given gene set. The gene sets in the top-most clade in the first branch of the row dendrogram are shown in the text box.

The gene sets associated with the most prominent of several clusters in Figure 7’s heatmap are delineated by the top-most clade in the first branch of the row dendrogram. Twenty gene sets in this clade have a relatively high occurrence among AS biclusters; 41% of the biclusters are enriched in at least one of these sets. Many of these sets include genes involved in antigenic variation, the export of cell adhesion proteins to the host cell membrane by the parasite, as well as erythrocyte-erythrocyte and erythrocyte-endothelium adherence. Each of these processes has been shown to be regulated epigenetically (64) and are central features of the parasite’s ability to evade immune system clearance. However, no single functional profile dominates among the AS biclusters. There are many additional functional gene sets associated with them, including - but not limited to - sets related to the cell cycle, metabolism, ribosome structural components, and ion transport. Thus, AS appears associated with coherence across a variety of biological functions, and by extension AR is associated with incoherence in these functions.

## Discussion

The EGRIN model of *P. falciparum* gene regulation described here is capable of making systems-level, quantitative predictions on samples from three separate validation data sets with a level of accuracy similar to those reported in previous EGRIN development efforts that were experimentally validated (19). We expect the model will serve the research community as a valuable hypothesis-generation tool for investigations into *P. falciparum’s* gene regulatory biology. The model’s predictive performance across validation data sets suggests that it is applicable to a variety of *P. falciparum* populations, including those clinically isolated from different geographic regions and from *in vitro* laboratory strains. Thus, it provides a critical step towards using transcriptomic data collected from field-isolated malaria parasites to predict critical parasite phenotypes and could contribute to tracking the emergence of drug resistance. Despite the heterogeneity in parasite origins and transcriptomic measurement techniques among the three validation data sets, the predictive performance of the model was generally consistent across validation tests. Even when tested against the third validation set, which differed from the training set in that it was obtained from an *in vitro* laboratory strain as opposed to clinical isolates and in that it consists of data from RNA-seq as opposed to microarray measurements, the model showed only a modest decrease in predictive performance (Figure 4).

Comparisons between model-based predictions of ApiAP2 TF targets and motif-based targets showed significant agreement for four out of 18 ApiAP2s tested. For the three TFs showing the highest level of agreement, mRNA abundances all peak during a relatively tight time interval during the early-mid schizont stage of the parasite’s intraerythrocytic developmental cycle (IDC) (8). Because our model uses gene expression coherence, as opposed to abundance, to identify co-regulated genes, the results of this analysis may be explained by timing differences between peak coherence of TF targets and peak abundance of TFs regulating those targets. Based on gene expression levels, the parasite populations used to train our model were found to be predominantly in the ring stage and expression levels of the top three ApiAP2s that showed significant agreement are relatively low (28). It may be that as the parasite initiates gene regulatory programs that transition the organism from stage to stage, ApiAP2-specific binding motifs become accessible and expression coherence among genes possessing those motifs is initially high. Then, as downstream regulatory mechanisms are activated by these programs, coherence decreases among those genes. In such a scenario, agreement between model-based and motif-based TF targets would be highest among TFs whose expression peaks in more downstream IDC stages. For TFs with peak expression during the ring stage, the model’s predicted targets would be influenced by the accumulation of systems-level changes in the parasite’s regulatory network brought on by increased TF expression, including feedback mechanisms influencing the expression of those TFs’ target genes. Thus, for our model-based versus motif-based target analysis, we do not necessarily expect agreement to be high among TFs with peak expression during the ring stage. We would instead expect agreement to be highest among TFs whose targets are newly transcriptionally active.

While the vast majority of predicted *P. falciparum* TRs require experimental validation, the model generated here can guide *P. falciparum* molecular biology research by identifying which proteins are likely to function as TRs, their predicted targets, binding sites and biological processes with which they associate. For example, by determining which TRs target genes are over-represented in biclusters enriched for AS samples, we have used the model to create a ranked list of putative TRs that appear to be associated with artemisinin sensitivity. We note that the TRs used in our model likely do not circumscribe the complete list of *P. falciparum* TRs. Nonetheless, the accuracy of our model indicates that the TR list we compiled is sufficient to generate a model with significant predictive power. While the model can potentially generate insights into *P. falciparum’s* gene regulatory programs, we may find that subsequent versions of the model with a refined list of TRs improves prediction accuracy and influences other model characteristics such as TR-target network topology.

Our network analysis of the EGRIN model suggests that the degree distribution of the *P. falciparum* gene regulatory network follows a power-law with a scaling exponent of 1.01, which is lower than those observed in networks derived empirically from studies on other model organisms. Previous research has found that for networks with scaling exponents less than 2, a relatively smaller set of nodes are required to control the entire network (65). Therefore, one interpretation of our results is that control of the *P. falciparum* gene regulatory network may be tighter compared to other organisms. However, additional post-transcriptional control of TRs, not incorporated into this model, likely also influences gene expression. Moreover, while our methodology quantifies the influence of each TR on all genes in the network, and the algorithm penalizes low-influence effects to exclude them from the network, the presence of relatively minor contributions of TRs to target gene expression may result in inflated TR out-degrees compared to the true physical biological network. This over-estimated connectivity could potentially result in lower power-law scaling exponents. Ultimately, a more comprehensive understanding of *P. falciparum* regulatory biology based on empirical studies is needed to validate the network characteristics derived from our model and to provide additional data to incorporate into future *Pf*EGRIN versions. We also note that the network characteristics derived from our model are based on data from blood stage parasites only. Whether these characteristics generalize across the various life cycle stages of the parasite requires further investigation. Moving forward, we are interested in directly comparing *P. falciparum’s* regulatory network characteristics to those of other organisms to illuminate the organism’s transcriptional control strategy and determine if and how the networks of obligate parasites such as *P. falciparum* differ from non-obligate organisms.

Overall, our efforts to find correlates of artemisinin resistance led to the discovery that resistance is linked to broad, increased transcriptional incoherence across a wide range of cellular processes (Figure 7). By comparison, hallmark constituents of the parasite’s epigenetically controlled mechanisms linked to virulence or the ability to persist under immune pressure (32, 66) appeared highly coherent in sensitive populations (Table S1, Figure 7). From these results, we hypothesize that artemisinin resistance may be an emergent property of parasite populations that have higher capacities for exploring diverse phenotypic states during infection. Parasite populations with more diversity in their gene regulatory programs may be better equipped to evade immune system clearance and establish infection sites, resulting in a higher parasite burden that requires more time to clear following artemisinin treatment. This observation also links immune evasion mechanisms with the capacity to circumvent drug pressure. This linkage, if substantiated, has critical consequences for malaria eradication, where current strategies include the use of mass drug administration, particularly in areas of high transmission that are populated with diverse, pre-existing anti-PfEMP1 immunity.

Bet-hedging, wherein organisms incur a fitness cost by generating phenotypically-diverse sub-populations that allow the overall population to persist in stressful conditions, is a proposed survival strategy of *P. falciparum* (31, 32). Our findings are consistent with parasite heterogeneity and bet-hedging being associated with survival in the presence of artemisinin. This is in contrast with the traditional view of the development of drug resistance, where it is presumed that a “sweep” of drug-sensitive parasites decreases genetic diversity within the population, yielding a relatively genetically homogenous set of parasites. A bet-hedging strategy has no requirement for genetic diversity. Rather, phenotypic and epigenetic heterogeneity, revealed by transcript incoherence present before drug treatment, enables a subset of the population to survive drug treatment. We hypothesize that the appearance of artemisinin resistance within a parasite infection may reflect a differentiation process analogous to the epigenetically-controlled transition from asexual to sexual parasite forms (66), allowing parasite populations to explore various phenotypic states. This process may be an inherent mechanism used by the parasite to keep blood stage populations sufficiently diverse so they persist under ongoing immune system stress.

Much of our current understanding of artemisinin resistance on a cellular level implicates macromolecular, global cellular processes. This is consistent with the notion that an assemblage of altered cellular states might precede resistance. These states include global processes such as production of phosphatidylinositol 3-phosphate containing vesicles (26), oxidative stress (27), the unfolded protein response (28) and hemoglobin endocytosis (29). Therefore, it would be reasonable to hypothesize that a systematic response within the parasite could modulate each of these cellular processes, leading to a diversity of cellular states, resulting in decreased artemisinin sensitivity. This is consistent with the extensive research on the function of the Kelch13 protein, which has been linked to some artemisinin resistant strains (24, 25) because various cellular pathways have been implicated in order to explain how Kelch13 contributes to resistance (29, 30). Within the context of the parasite’s bet-hedging mechanisms, investigations into whether Kelch13 modulates the phenotypic diversity of clonal parasite populations, and whether mutations in the protein increase that diversity, would be warranted.

Our results illustrate a key advantage in using systems-level perspectives to understand infectious disease processes such as drug resistance. The holistic, integrated perspective we have applied here has illuminated widespread differences in gene expression coherence between AS and AR infections. Based on our functional profiling results (Figure 7), there is a clear collection of biological functions associated with AS biclusters that involve genes contributing to antigenic variation and erythrocyte-host interactions. These associations are driven largely (but not exclusively) by the over-representation of *var* and *rifin* genes among a subset of the AS biclusters. We acknowledge that interpreting microarray results that focus on *var* and *rifin* genes can be problematic, given the high sequence complexity in these genes and the potential cross-reactivity among microarray probes that target them. Thus, we hesitate to draw conclusions that implicate specific *var* or *rifin* genes as correlates of AR. We also emphasize that most AS biclusters do not contain members of these gene families and are associated with other cellular processes. The strong association observed between artemisinin sensitivity and genes responsible for antigenic variation and erythrocyte-host interactions may be a result of those processes being some of the most well-studied in *P. falciparum*, given their clinical relevance. In contrast, many *P. falciparum* proteins remain functionally uncharacterized and there may be critical, undiscovered gene regulatory pathways that cannot yet be revealed by functional enrichment analysis and thus remain occult in our assessment of the functional landscape of AS biclusters. AS biclusters that show more fragmented functional profiles may in fact represent key gene regulatory programs in the parasite that have yet to be characterized and aggregated into broader functional categories. The biclusters we generated for this study, because they implicate proteins in related functions, can offer a potential guide for functionally characterizing the parasite’s less-studied proteins. Future research focused on characterizing these proteins and the pathways in which they participate might substantially benefit from our *Pf*EGRIN model, as it provides a collection of hypotheses about which proteins participate in and/or regulate critical transcriptional functions.

## Acknowledgements

We thank Robert Morrison for guidance on the use of his software for functionally profiling *P. falciparum* gene sets. We thank Mari Couasnon for exploratory work that helped initiate the *P. falciparum* EGRIN modeling project. We also thank Joseph D. Smith and Jason Wendler for helpful discussions related to artemisinin resistance and parasite heterogeneity.

