## Supplementary material for "A system-level gene regulatory network model for *Plasmodium falciparum*": EGRINtools.R

```
# Get location of script
getScriptDir <- function(){
  return(dirname(sys.frame(1)$ofile))
}

# Step through samples and make predictions
predictGeneAcrossSamples <- function(TRexp=NA, edgedata=NA, genei=NA, predVals=NA){
  for(samplei in 2:dim(TRexp)[2]){
    
    TRvals <- TRexp[,samplei]
    names(TRvals) <- TRexp[,"Geneid"]
    weights <- edgedata$weight
    
    # Get the predicted value for the gene's relative expression
    predVals[genei,samplei] <- predictExpValue(TRvals, weights, names(TRexp)[samplei])
  }
  
  return(predVals)
}


# From expression levels of TRs that influence a gene, and the Inferelator-based weight, make prediction on new data
predictExpValue <- function(TRlevels=NA, weights=NA, sample=NA){
  predVal <- 0
  for(i in 1:length(TRlevels)){
    predVal <- predVal + (TRlevels[i] * weights[i])
  }
  return(predVal)
}


# Get the genes and samples for a bicluster
getCachedBiclusterGenes <- function(cache=NA, runi=NA, clust=NA){
  
  geneandsampleset <- list()
  
  message(c("Looking up conditions and genes for run ", runi, " and bicluster ",clust))
  # lookup cluster at last iteration and get its row members and column members
  condnames <- strsplit(cache[cache$RunNumber==runi & cache$BiclusterNumber==clust,"Samples"],split = ';')[[1]]
  rownames <- strsplit(cache[cache$RunNumber==runi & cache$BiclusterNumber==clust,"Genes"],split = ';')[[1]]
  
  geneandsampleset[[1]] <- rownames
  geneandsampleset[[2]] <- condnames
  
  names(geneandsampleset) <- rep(paste0("Run ", runi, " Bicluster ", clust),2)
  
  return(geneandsampleset)
}
```
