## Supplementary material for "A system-level gene regulatory network model for *Plasmodium falciparum*": makePredictionOnExampleBicluster.R

```
# - - - - - - - - - - - - - - - - - - - - - - - - - - - - - - - - - - - - - - - - #
# Run this script as source or as R script and it will output an image that 
# reproduces the EGRIN model prediction results for a representative bicluster from
# validation set 1 (GSE59098)
# - - - - - - - - - - - - - - - - - - - - - - - - - - - - - - - - - - - - - - - - #


# Function to figure out the location of this file on the user's machine
thisFile <- function() {
  cmdArgs <- commandArgs(trailingOnly = FALSE)
  needle <- "--file="
  match <- grep(needle, cmdArgs)
  if (length(match) > 0) {
    # Rscript
    return(normalizePath(sub(needle, "", cmdArgs[match])))
  } else {
    # 'source'd via R console
    return(normalizePath(sys.frames()[[1]]$ofile))
  }
}

locdir <- thisFile()
locdir <- gsub("makePredictionOnExampleBicluster.R$","",locdir)

source(paste0(locdir, "EGRINtools.R")) # Load functions used to make predictions

message("Loading model...")
load(paste0(locdir, "EGRIN.RData")) # Loads the TR-target interaction network

message("Loading test data...")
load(paste0(locdir, "TestData.RData")) # Loads validation data set 1

message("Loading training data...")
load(paste0(locdir, "TrainingData.RData")) # Loads the training data

message("Loading example bicluster...") # Loads the example bicluster
load(paste0(locdir, "ExampleBicluster.RData"))

geneandsampleset <- getCachedBiclusterGenes(cache=ExampleBicluster, 1, 28)
rownames <- unlist(geneandsampleset[1])
condnames <- unlist(geneandsampleset[2])

actualValsTest <- TestData[TestData$Geneid %in% rownames,] # Get actual gene expression values from test set
actualValsTest <- actualValsTest[order(actualValsTest$Geneid),]

ratiosTrainInBi <- TrainingData[ , c("Geneid",condnames)]

actualValsTrainInBi <-ratiosTrainInBi[ratiosTrainInBi$Geneid %in% rownames,] # Get actual gene expression values from samples in training set that are in bicluster
actualValsTrainInBi <- actualValsTrainInBi[order(actualValsTrainInBi$Geneid),] 

predValsTest <- matrix(nrow=dim(actualValsTest)[1], ncol=dim(actualValsTest)[2]) # Set up data frame that will hold predictions for test data
predValsTest <- as.data.frame(predValsTest)
names(predValsTest) <- names(actualValsTest)
predValsTest$Geneid <- actualValsTest$Geneid

predValsTrainInBi <- matrix(nrow=dim(actualValsTrainInBi)[1], ncol=dim(actualValsTrainInBi)[2]) # Set up data frame that will hold predictions for training data (samples in bicluster)
predValsTrainInBi <- as.data.frame(predValsTrainInBi)
names(predValsTrainInBi) <- names(actualValsTrainInBi)
predValsTrainInBi$Geneid <- actualValsTrainInBi$Geneid

tempedges <- EGRIN[EGRIN$Target %in% rownames,]
TRexpTest <- TestData[TestData$Geneid %in% tempedges$TR,]
TRexpTest <- TRexpTest[order(TRexpTest$Geneid),]

# For each gene in bicluster make prediction
for(genei in 1:dim(actualValsTest)[1]){
  
  geneid <- actualValsTest$Geneid[genei]
  
  # collect all TRs that influence the gene's expression
  edgedata <- EGRIN[EGRIN$Target==geneid,]
  edgedata$weight <- edgedata$weight
  edgedata <- edgedata[order(edgedata$TR),]
  
  # Get the TR expression levels from the test data
  TRexpTest <- TestData[TestData$Geneid %in% edgedata$TR,]
  TRexpTest <- TRexpTest[order(TRexpTest$Geneid),]
  
  TRexpTrainInBi <- ratiosTrainInBi[ratiosTrainInBi$Geneid %in% edgedata$TR,]
  TRexpTrainInBi <- TRexpTrainInBi[order(TRexpTrainInBi$Geneid),]
  
  predValsTest <- predictGeneAcrossSamples(TRexpTest, edgedata, genei, predValsTest)
  predValsTrainInBi <- predictGeneAcrossSamples(TRexpTrainInBi, edgedata, genei, predValsTrainInBi)
}

actualMeansTest <- colMeans(actualValsTest[,-1], na.rm = F)
predMeansTest <- colMeans(predValsTest[,-1], na.rm = F) 

actualMeansTrainInBi <- colMeans(actualValsTrainInBi[,-1], na.rm = F)
predMeansTrainInBi <- colMeans(predValsTrainInBi[,-1], na.rm = F) 

OKsamplesiTest <- intersect(which(!is.na(predMeansTest)), which(!is.na(actualMeansTest)))
predMeansTest <- predMeansTest[OKsamplesiTest]
actualMeansTest <- actualMeansTest[OKsamplesiTest]

OKsamplesiTrainInBi <- intersect(which(!is.na(predMeansTrainInBi)), which(!is.na(actualMeansTrainInBi)))
predMeansTrainInBi <- predMeansTrainInBi[OKsamplesiTrainInBi]
actualMeansTrainInBi <- actualMeansTrainInBi[OKsamplesiTrainInBi]
  
RMSEtest <- (mean((predMeansTest - actualMeansTest)^2, na.rm=T)^0.5)
RMSEtrainInBi <- (mean((predMeansTrainInBi - actualMeansTrainInBi)^2, na.rm=T)^0.5)

corTest <- cor.test(actualMeansTest,predMeansTest)
corTrainInBi <- cor.test(actualMeansTrainInBi,predMeansTrainInBi)


# Generate output image
title <- paste0("Measured (red) and predicted (blue) gene expression")
outputfile <- paste0(locdir, "ExampleBiclusterPredictions.png")

png(filename=outputfile, width=18, height=10.5, units="in", res=300)

par(mfrow=c(2,2))
par(mar = c(8, 10, 6, 6)) # b, l, t, r

plot(0,0, cex.main = 1.7, cex.lab=2, cex.axis=2, xaxt="n", ylim=c(-4,4),xlim=c(0,length(predMeansTest)-1),
     type = "n", main=paste0(title, "\nRMSE test data: ", round(RMSEtest, digits=2)), 
     xlab="Test data samples", ylab="Normalized\ngene expression")
lines(0:(length(predMeansTest)-1), actualMeansTest, col = "red", type = 'l', lwd=1.3)
lines(0:(length(predMeansTest)-1), predMeansTest, col = rgb(0,0,1), type = 'l')
axis(1, at=0:(length(predMeansTest)-1), labels=rep("",length(predMeansTest))) 

plot(x=actualMeansTest, y=predMeansTest, cex.main = 1.7, cex.lab=2, cex.axis=2, cex=2, 
     main=paste0("Correlation R: ", round(corTest$estimate, digits=2),"     Correlation P-value: ", 
                 round(corTest$p.value, digits=10)), xlab="Measured value", ylab="Predicted value")

plot(0,0, cex.main = 1.7, cex.lab=2, cex.axis=2, xaxt="n", ylim=c(-4,4),xlim=c(0,length(predMeansTrainInBi)-1),
     type = "n", main=paste0(title, "\nRMSE training data (in bicluster): ", round(RMSEtrainInBi, digits=2)),
     xlab="Training data samples", ylab="Normalized\ngene expression")
lines(0:(length(predMeansTrainInBi)-1), actualMeansTrainInBi, col = "red", type = 'l', lwd=1.3)
lines(0:(length(predMeansTrainInBi)-1), predMeansTrainInBi, col = rgb(0,0,1,0.8), type = 'l')
axis(1, at=0:(length(predMeansTrainInBi)-1), labels=rep("",length(predMeansTrainInBi)))

plot(x=actualMeansTrainInBi, y=predMeansTrainInBi, cex.main = 1.7, cex.lab=2, cex.axis=2, cex=2, 
     main=paste0("Correlation R: ", round(corTrainInBi$estimate, digits=2),"     Correlation P-value: ", 
                 round(corTrainInBi$p.value, digits=10)), xlab="Measured value", ylab="Predicted value")

dev.off()

message("Output image written to ", outputfile)
```
