## Supplementary material for "A system-level gene regulatory network model for *Plasmodium falciparum*": README.txt

This directory contains files for reproducing P. falciparum EGRIN 
model predictions on test data from GEO accession GSE59098.

Running the script makePredictionOnExampleBicluster.R will generate
a PNG image that reproduces model prediction results for genes
in an example bicluster.

Contact: max [dot] neal [at] seattlechildrens [dot] org
