## Supplementary material for "A system-level gene regulatory network model for *Plasmodium falciparum*": Figure S1

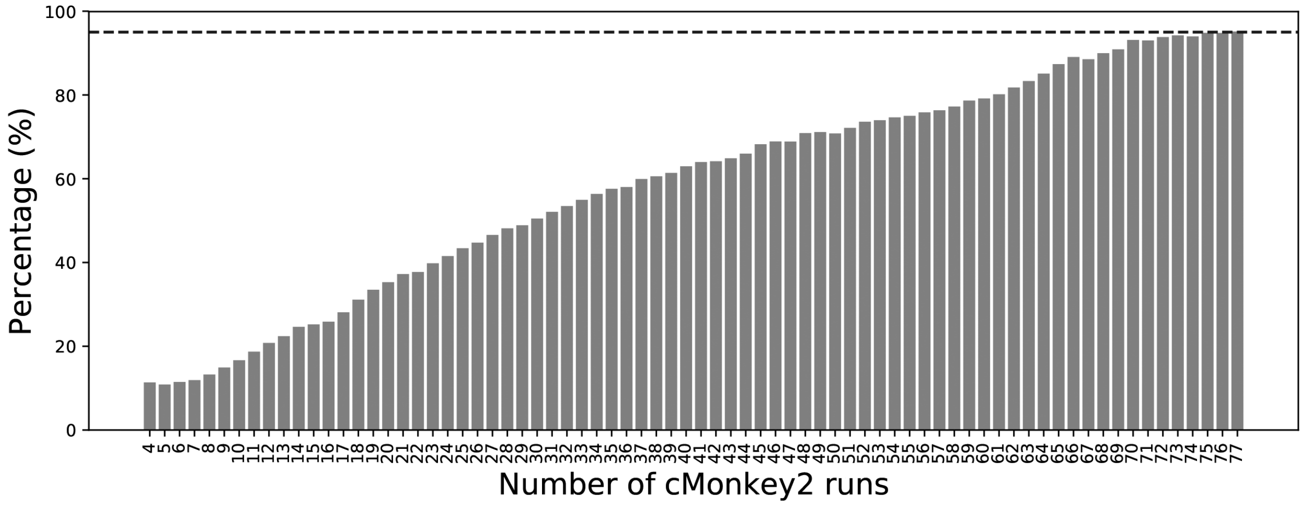


**Figure S1.** Convergence of average TR-target weights over 77 cMonkey2/Inferelator runs. Bars indicate the percentage of TR-target pairs with weights that differed less than 5% between *n* and *n*-3 cMonkey2 runs, with *n* given on the x-axis.
