## Supplementary figures and images for "A system-level gene regulatory network model for *Plasmodium falciparum*"

### Figure S2

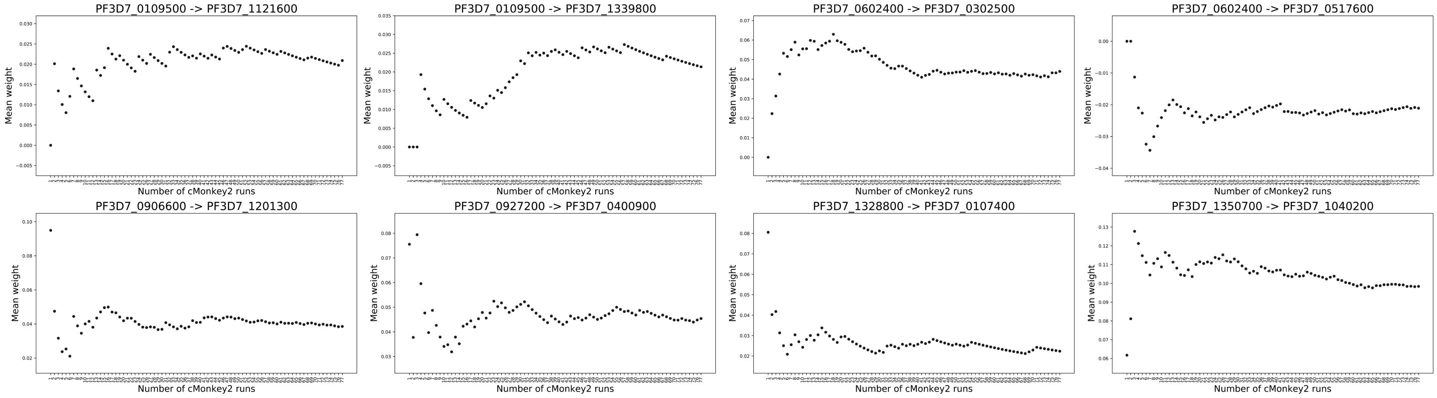


**Figure S2.** Examples of specific mean TR-target weights over 77 cMonkey2/Inferelator runs.
