## Supplementary material for "A system-level gene regulatory network model for *Plasmodium falciparum*": Figure S3

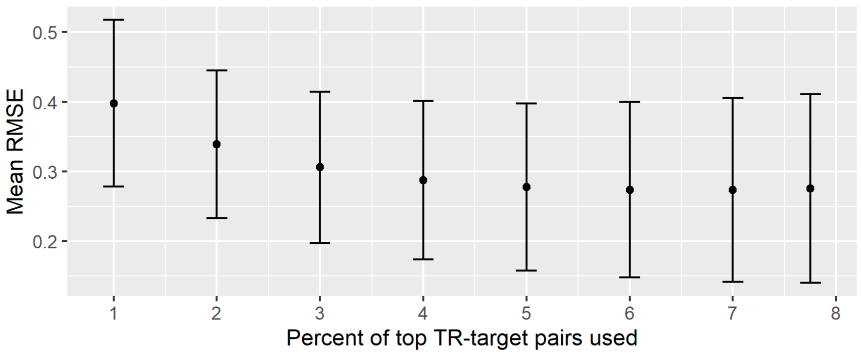


**Figure S3.** Mean RMSE values for model fits to training data versus percent of top-weighted TR-target pairs used in the *Pf*EGRIN model. Model fits were assessed using gene sets from the 38,500 biclusters generated to build the model. An optimal RMSE value was found using the top 7% of pairs.
