## Supplementary material for "A system-level gene regulatory network model for *Plasmodium falciparum*": Figure S4

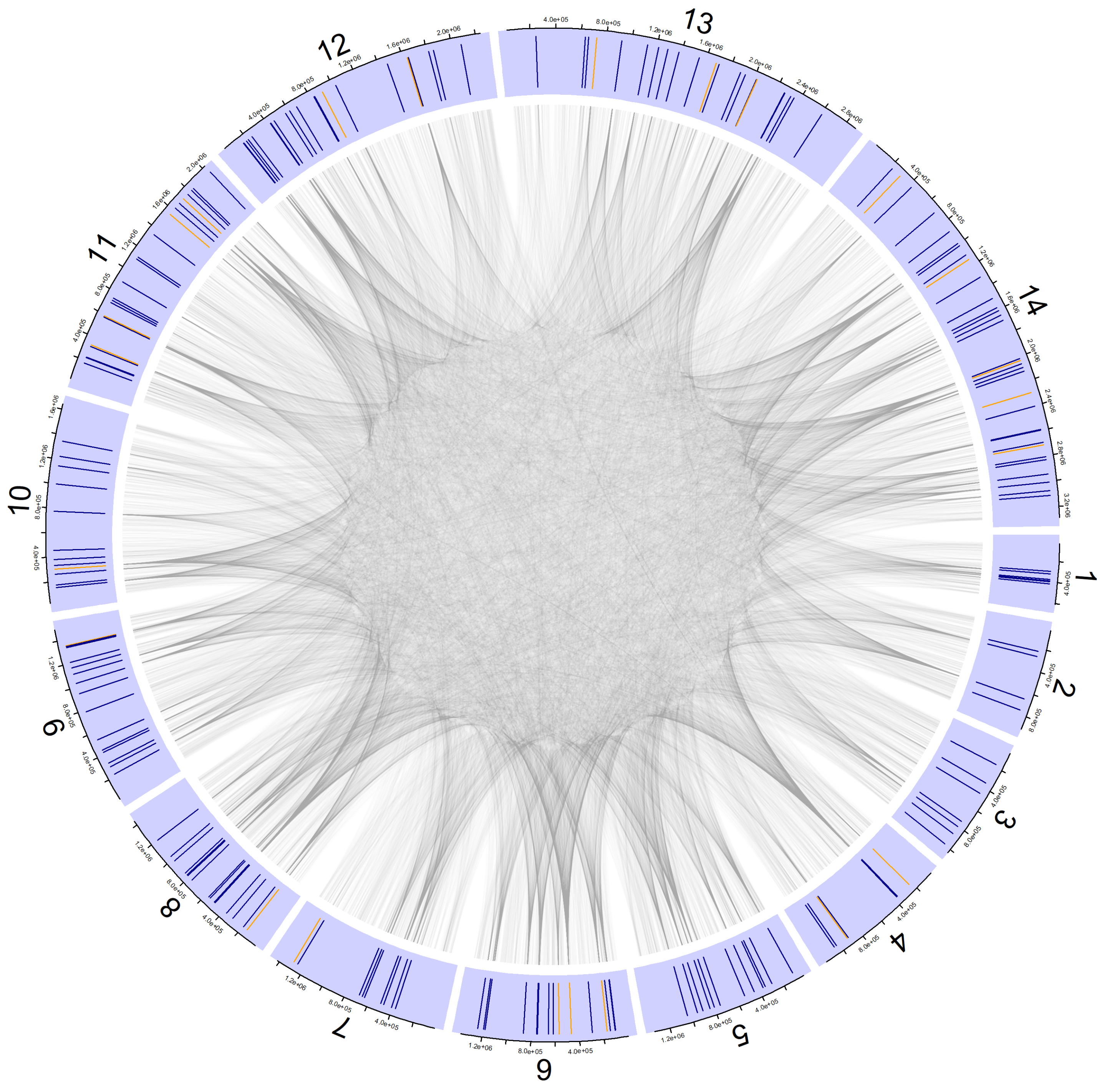

**Figure S4.** Visualization of gene regulatory interactions across the *P. falciparum* genome. Segments of the ring represent *P. falciparum* chromosomes. Lines inside segments indicate genomic location of TRs used in the *Pf*EGRIN model. Orange lines indicate ApiAP2 genes; blue lines indicate other TRs. Edges indicate TR-target interactions positioned according to the genomic location of interacting genes. For visualization purposes, only edges with absolute weights within the top quartile are shown.
